# Accurate and memory-efficient cell type annotation from multimodal single-cell RNA and protein data

**DOI:** 10.64898/2026.04.24.717187

**Authors:** Siddharth S. Tomar, Joel T. Haas, Bart Staels, David Dombrowicz, Jonas Nørskov Søndergaard

## Abstract

Accurate cell annotation remains a central challenge in single-cell analysis, particularly when datasets contain rare populations, transitional states, and tissue-adapted phenotypes. We previously developed scODIN as an expert-guided framework for immune cell annotation in single-cell RNA-sequencing data. Here, we present pyODIN, a major extension of this framework for Python. pyODIN supports multimodal annotation using RNA, antibody-derived tag (ADT), or combined RNA-ADT information. It also substantially expands the annotation database from a CD4 T-cell-centred framework to a broader cell-type reference resource spanning major and minor peripheral blood cell populations as well as tissue-associated subsets. We benchmarked pyODIN against CellTypist, MMoCHi and HiCAT using independently curated immune reference datasets. pyODIN demonstrated the highest overall classification accuracy, and at the top-lineage level, pyODIN reduced cross-lineage errors relative to the benchmark methods. At finer resolution, pyODIN preserved substantially greater CD4 T-cell subtype structure than CellTypist and HiCAT, resolving regulatory, follicular, helper, memory, and cytotoxic states that were collapsed into broader categories by the comparator methods. In a controlled marker-ablation experiment, pyODIN outperformed MMoCHi when key lineage-defining RNA markers were removed from the expression feature set, and the addition of ADT information restored NK and CD8 T-cell annotation, demonstrating the value of multimodal annotation when transcript-level evidence is incomplete. Finally, application to liver fine-needle aspirate data showed that the expanded framework supports annotation beyond PBMC datasets. Together, pyODIN provides an expert-guided, adaptable cell annotation framework for Python, designed for multimodal cell phenotyping across blood and tissue single-cell datasets. pyODIN is freely available at https://github.com/SondergaardLab/pyODIN.

## Introduction

Single-cell transcriptomics has transformed immunology by enabling high-resolution profiling of immune heterogeneity across homeostasis, infection, autoimmunity, metabolic disease and cancer[1–3]. However, biological insight depends critically on accurate cell-type annotation. This remains challenging in practice because immune populations are often separated by only a limited number of lineage-defining markers, more abundant cell types often obscure rare populations, and transcript dropout can mask key genes required for confident assignment[4, 5]. These challenges become even greater when analyses move beyond broad lineage classification toward fine-grained immune subset annotation.

A growing number of computational methods have been developed for automated cell annotation. These include reference-based transcriptome classifiers as well as multimodal approaches for CITE-seq data[6, 7]. While such tools have substantially improved scalability, several limitations remain particularly relevant for immunology. First, performance on rare or closely related immune populations often remains weaker than performance on abundant populations[8]. Second, many tools are relatively static from the perspective of biological curation, making it less straightforward to update subtype definitions as immune knowledge evolves. Third, transitional, mixed, and tissue-adapted immune phenotypes are often difficult to represent in a biologically interpretable manner[9]. Finally, multimodal support remains limited across various methods and datasets, despite its potential to facilitate better separation of immune cell populations.

We previously introduced ”Optimized Detection and Inference of Names in scRNA-seq data” (scODIN) as an expert-guided annotation framework that combined biologically curated marker priorities with machine-learning-based inference of labels to incompletely resolved cells[10]. The original implementation demonstrated the value of informed immune annotation logic and was developed primarily around detailed CD4 T-cell phenotyping. In addition, scODIN was explicitly designed to identify mixed and intermediate phenotypes and to extend confident core annotations to less certain cells through a machine-learning-based inference step. Although this framework proved useful, its detailed database scope initially focused on CD4 T-cell biology, and its implementation was tied to the R ecosystem.

Here we present pyODIN, a Python implementation of scODIN that supports annotation from RNA alone, antibody-derived tag (ADT) alone, or combined RNA-ADT information, enabling direct application to both conventional scRNA-seq and multimodal single-cell datasets[7, 11]. This is particularly useful for resolving immune populations that can be difficult to distinguish robustly in transcriptomic data, including CD4 T cells, CD8 T cells, NK cells, and γδ T cells, especially when lineage-defining markers are weakly detected[7]. In parallel, the underlying annotation resource has been expanded from a CD4+ T cell-focused framework to a broader immune phenotyping database spanning major and minor blood immune populations, including CD8+ T cells, B cells, dendritic cells (DCs), and monocytes, as well as liver tissue-resident immune subsets such as CD8+ tissue-resident memory (TRM) cells, monocyte-derived and resident macrophages, including Kupffer cells[9, 12–14]. Conceptually, pyODIN complements existing annotation strategies by combining an editable, expert-guided immune knowledge database with multimodal support and explicit accommodation of mixed or transitional phenotypes. While CellTypist[15] relies on pretrained references, HiCAT[16] relies on database-derived RNA models, and MMoCHi[17] relies on a custom, user-defined cell-type identification hierarchy, pyODIN distinguishes itself by incorporating curated immunological expertise directly into the annotation workflow.

In this study, we present the pyODIN framework and benchmark its performance across independent immune reference atlases, demonstrating improved annotation performance compared with the benchmark methods tested here.

## Methods

pyODIN is based on the scODIN framework, which combines expert-curated marker knowledge with machine-learning-based inference to assign cell identities in large single-cell datasets [10]. pyODIN shares the core architecture of scODIN, with the Seurat framework replaced by the Scanpy/AnnData ecosystem in Python. The gene priority table format is interchangeable between the two tools.

### Data architecture and multimodal processing

pyODIN operates on single-cell datasets stored in the AnnData format and supports simultaneous analysis of transcriptomic (RNA) and proteomic (antibody-derived tag, ADT/CITE-seq) data. RNA features are read from either the primary expression matrix (.X) or a specified layer (e.g. scaled data), falling back to .X if no layer is specified or the requested layer is absent; only the (typically much smaller) subset of genes listed in the marker table is sliced out of the expression matrix before any sparse-to-dense conversion, keeping memory use low even for very large gene expression matrices (see *Computational implementation*). ADT data are read from the obsm slot in full; since ADT panels are comparatively low-dimensional (typically tens of antibodies) and are conventionally stored as a dense array or DataFrame in obsm, no comparable subsetting or densification step is required for this modality - only the antibody columns referenced by the marker table are used once scores are computed.

### Weighted scoring framework

Each cell type is defined in a curated marker table by a set of marker genes and/or ADT targets, each with an assigned weight and tier. A marker gene shared by more than one cell type (e.g. *CD3D* for both T cells and NKT cells) is scored independently for each type it is associated with, using that type’s own weight. Only markers actually present in the dataset are used, so the effective marker set for each cell type can vary between datasets.

For a given cell type *c*, the per-cell score is the weighted sum of its marker values, normalised by the square root of the number of markers used for that cell type:

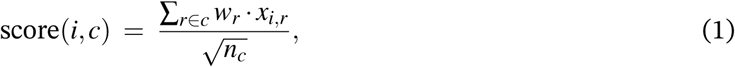

where *x_i,r_* is the expression of marker *r* in cell *i*, *w_r_* is that marker’s curated weight, the sum runs over all markers *r* assigned to cell type *c*, and *n_c_* is the number of such markers. This normalisation prevents cell types defined by many markers from being unfairly favoured over those defined by only a few. When both RNA and ADT are used, *x_i,r_* is simply the RNA value, the ADT value, or their sum, depending on which modality is active for that marker; by default, ADT markers are only counted when annotated as a positive marker for the cell type (negative ADT markers are ignored unless this default is disabled), while RNA markers are used regardless of their annotated direction. This is due to higher background of the ADT signal. Scores below a user-defined core-cell cutoff (default 5.0) are set to zero, so that only cell types with reasonably strong marker support retain a non-zero score for a given cell.

For computational efficiency, the scoring can be described as *X* ∈ ℝ*^N^*^×*F*^ for the matrix of active marker values across all *N* cells and *F* markers, and *W* ∈ ℝ*^F^*^×*K*^ for the marker-weight matrix (with *W_r,c_* = *w_r_* if marker *r* is assigned to cell type *c*, and 0 otherwise), the same computation is a single matrix product,

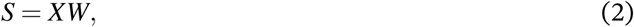

with each column of *S* subsequently divided by 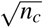 and thresholded at the core-cell cutoff. This matrix formulation is what enables the vectorised implementation described below.

### Hierarchical tiering and labelling logic

Cell types are grouped into ordered tiers, so that rare cells are identified first before common cell types, which may share markers with the rare cell types. Labelling proceeds tier by tier: at each tier, only cells still unlabelled from previous tiers are considered. For each such cell *i* and tier *t*, let *v*_1_ and *v*_2_ denote the highest and second-highest score among the cell types in tier *t*:

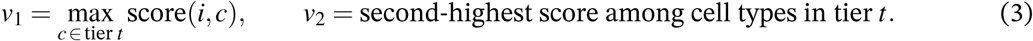

The cell is then labelled according to the following rule:

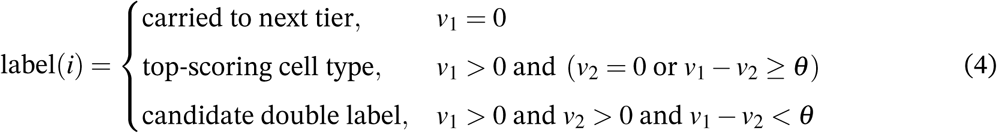

where θ is a similarity threshold (default 1.0). Note that an exact tie between the top two scores (*v*_1_ = *v*_2_ *>* 0) falls under the double-label case, since a difference of zero is still below any positive threshold.

Candidate double labels are checked against a curated list of biologically accepted double labels (e.g. transitional or mixed phenotypes). If the pair is on this list, the cell receives the accepted double label, which is then mapped to a simplified, human-readable name. If the pair is not on the list, the outcome depends on which data modality was used: for RNA-only annotation, the cell is still given the higher-scoring of the two types as a provisional label; for ADT-only or combined RNA+ADT annotation, the cell is instead left labelled unknown, reflecting lower confidence in resolving the ambiguity from a single line of evidence when a second modality was available but inconclusive. Cells that score zero for every cell type across all tiers are likewise labelled unknown. All unknown cells are candidates for the machine-learning-based label propagation step described below.

### Cluster-based consensus scoring

To reduce the impact of cell-level noise, per-cell scores can be summarised at the level of a predefined clustering (e.g. Leiden clusters). Let κ(*i*) denote the cluster assignment of cell *i*. For cluster *k* and cell type *c*, the cluster-level total score is the sum of per-cell scores over all cells in that cluster:

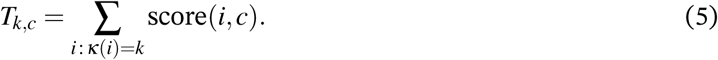

To keep this step computationally efficient on large datasets, candidate cell types for a cluster are restricted to those that are *both* assigned to at least one cell in that cluster at the single-cell level *and* rank among the ten cell types with the largest *T_k,c_* within that cluster (all cell types are eligible on the score criterion if there are fewer than ten in total).

For computational efficiency, writing *C* ∈ {0, 1}*^n^*^clusters×^*^N^* for the cluster-membership matrix (*C_k,i_* = 1 if cell *i* belongs to cluster *k*, and 0 otherwise) and *S* for the per-cell score matrix (Eq. 2), the same aggregation is a single matrix product,

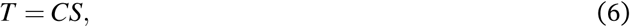

with *T_k,c_* the (*k, c*) entry of *T* .

For each remaining candidate cell type, a ranking score is computed by combining its cluster-level total score with *n_k,c_*, the number of cells in cluster *k* labelled *c* at the single-cell level:

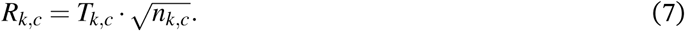

The cell type with the largest *R_k,c_* becomes the cluster’s consensus label, 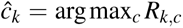. As a diagnostic, an enrichment ratio *E_k_* = *R_k,c_*(1) */R_k,c_*(2) – the ranking score of the top candidate divided by that of the runner-up - is reported alongside each cluster’s consensus label, indicating how clearly the winning label separates from the next-best alternative; this ratio is not itself used to decide the assignment. If cluster-based consensus is used, we recommend over-clustering with a high resolution (>2). Cluster-based consensus is, however, not recommended for annotation at the subtype level.

### pyODIN ambiguous cell classification

Cells labelled unknown due to ambiguity or marker dropout are resolved using a supervised gradient-boosted classifier (LightGBM) [18], in place of the kNN-based label transfer (FindTransferAnchors/ TransferData) used in the original scODIN implementation.

Before training, cell types represented by only a single cell are excluded, since they would prevent stratified cross-validation. The remaining labelled cells are split 80/20 into a training set and a validation set (stratified by cell type, fixed random seed), and sample weights inversely proportional to each class’s frequency are applied throughout training to counteract class imbalance between common and rare cell types.

A multiclass LightGBM model is first trained on the 80% training split, using a feature fraction of 0.2 (each tree considers a random 20% of features, which helps regularise the model given the high dimensionality of the gene expression matrix), a learning rate of 0.03, 31 leaves per tree, and a minimum of 5 samples per leaf. The 20% validation split is used to monitor multiclass log-loss, with training stopped early if no improvement occurs for 30 consecutive boosting rounds (up to a maximum of 1000 rounds), identifying the optimal number of boosting rounds. A final production model is then retrained with this optimal number of rounds on all labelled cells combined, so that the classifier used for prediction benefits from the full reference set.

For each unknown cell *i*, the production model outputs a predicted probability *p_i,k_* for every reference cell type *k*. The predicted label is *y*^*_i_* = argmax*_k_ p_i,k_*, with confidence max*_k_ p_i,k_*. Cells are assigned *y*^*_i_* directly if max*_k_ p_i,k_* ≥ 0.65 (default); otherwise they are labelled Uncertain_*y*^*_i_*, preserving the model’s best guess. The feature matrix used for training and prediction was the normalized and scaled gene count matrix (adata.X).

### Python implementation

To scale to large single-cell datasets, per-cell scoring, tiered labelling, and cluster-level aggregation are implemented as vectorised array operations rather than explicit loops over cells or cell types. Per-cell scoring is computed as a single dense matrix multiplication, *S* = *XW* (Eq. 2), using NumPy routines in place of a Python-level loop over cell types. Sparse RNA inputs are densified only over the marker-derived gene subset actually required for scoring (typically tens to a few hundred genes), substantially reducing peak memory relative to densifying the full transcriptome; this optimisation is specific to RNA, as ADT panels are small and already stored densely. Within tiered labelling, the top two candidate scores per cell are identified via partial selection (argpartition) rather than a full sort, avoiding the *O*(*K* log *K*) cost of sorting all cell-type scores per cell when only the top two are required. Cluster-level aggregation uses a sparse cell-to-cluster indicator matrix and a single sparse–dense matrix product, *T* = *CS* (Eq. 6), in place of an explicit loop over clusters. Exact versions of the tools and libraries used, along with the workstation configuration, are provided in Table S1.

### Comparison to other algorithms

pyODIN was benchmarked against three established annotation tools, HiCAT, CellTypist and MMoCHi[15–17] using curated single-cell datasets from the Allen Institute and Azimuth references [7, 11]. Both datasets underwent quality control and preprocessing with an identical set of parameters. The Immune_All_Low model was used for CellTypist, and the HiCAT-supplied database (cell_markers_rndsystems_hs.tsv) from the HiCAT repository was used for HiCAT. All tools were run with default configurations, following the tutorials provided by their respective authors. For preliminary analysis, cell-type labels were collapsed and standardised across tool predictions and author annotations to ensure direct comparability (Table S2); for example, “Transitional B cells” and “Treg(diff)” were mapped to the broader categories “B cell” and “CD4^+^ T cell”, respectively. For MMoCHi, we used the example hierarchy (Human immune subsets (v1)) provided in the tool documentation. We substituted the ADT tag names with the corresponding ones in the Azimuth dataset (which contains multiple antibodies with custom names).

### Application to liver fine-needle aspirate data

pyODIN was applied to a published human liver fine-needle aspirate single-cell RNA-seq dataset spanning the MASLD/MASH spectrum (dbGaP accession phs004044.v1.p1) [3] to assess annotation performance in a disease-relevant tissue context.

## Results

### pyODIN extends the ODIN framework into a multimodal and cross-compartment annotation platform

We developed pyODIN as a Python-native implementation of the ODIN framework for single-cell annotation. Relative to the original scODIN implementation, pyODIN introduces three major advances. First, it enables ODIN-based annotation within Python-centered workflows built around Scanpy and AnnData (Fig. 1a). Second, it substantially broadens the underlying annotation database from a CD4 T-cell-focused resource to a wider annotation framework spanning major PBMC immune populations together with tissue-associated immune and non-immune cell states (Fig. 1b-d). Third, it extends the framework from transcriptome-only annotation to multimodal annotation using RNA, ADT, or combined RNA+ADT evidence (Fig. 1e). In addition, pyODIN retains the expert-curated and tier-aware logic of the original ODIN framework and machine-learning-based propagation of confident labels to unresolved cells (Fig. 1f). Whereas the original scODIN framework was primarily developed for detailed CD4 T-cell phenotyping, pyODIN is designed as a broader platform for both top-level and fine-grained annotation across blood and tissue datasets. At the PBMC level, the framework supports detailed phenotyping of CD4 T cells, CD8 T cells, B cells, myeloid/DC populations, and NK-cell subsets (Fig. 1c). At the tissue level, the database has been extended to include liver-associated macrophages, CD8 tissue-resident memory (TRM) phenotypes, and broad non-immune tissue categories (Fig. 1d).

**Figure 1:**
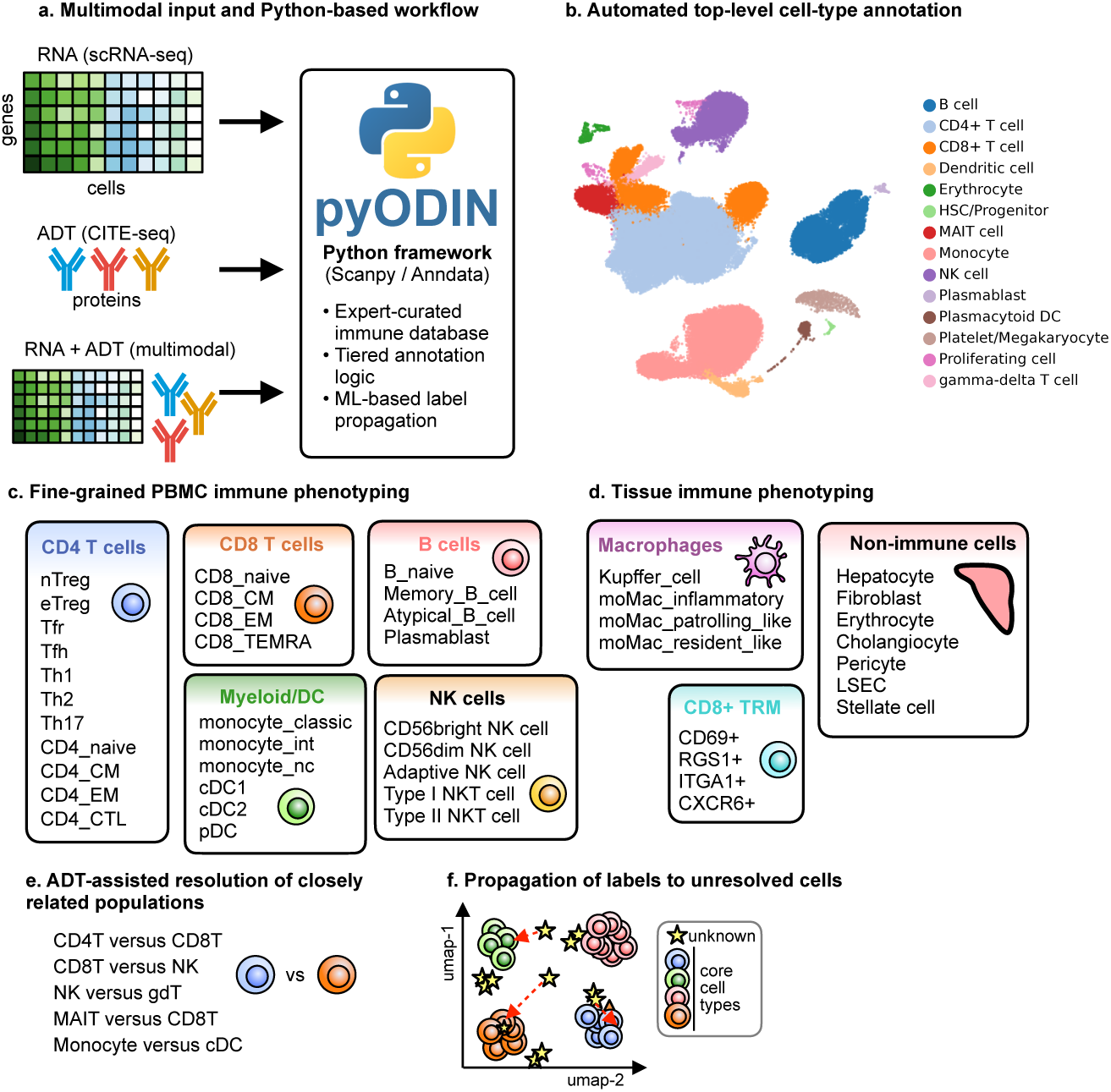
Overview of the pyODIN framework for multimodal single-cell annotation. **(a)** pyODIN is a Python-based framework implemented for Scanpy/AnnData workflows and can operate on RNA (scRNA-seq), antibody-derived tag (ADT; CITE-seq), or combined RNA+ADT input. The framework combines an expert-curated immune annotation database with tiered annotation logic and machine-learning-based label propagation. **(b)** At the top level, pyODIN performs automated broad cell-type annotation across major immune cell populations. **(c)** pyODIN enables fine-grained PBMC immune phenotyping across multiple compartments, including detailed annotation of CD4 T-cell, CD8 T-cell, B-cell, myeloid/DC, and NK-cell subsets. **(d)** The database also supports tissue immune phenotyping, illustrated here for liver-associated cell states including macrophage subsets, CD8+ tissue-resident memory (TRM) cells, and broad non-immune tissue cell categories. **(e)** Multimodal annotation using ADT information can assist the resolution of closely related populations that are difficult to distinguish robustly from transcriptomic information alone. **(f)** Confidently annotated core cells are used to propagate labels to unresolved cells with weak or ambiguous signals.

### pyODIN shows strong performance for top-level immune cell annotation

We first benchmarked pyODIN against CellTypist and HiCAT for top-level annotation using two independently curated immune reference atlases, referred to here as the Allen[11] and Azimuth[7] datasets. pyODIN achieved the highest overall classification accuracy on the Allen dataset and matched the top-performing methods on Azimuth (Fig. 2a). Specifically, pyODIN reached accuracies of 0.99 and 0.91 on the Allen and Azimuth datasets, respectively, compared with 0.96 and 0.92 for CellTypist. HiCAT could not be run on the Allen dataset due to excessive memory usage (*>* 512GB), but achieved an accuracy of 0.51 on Azimuth (Fig. 2a).

**Figure 2:**
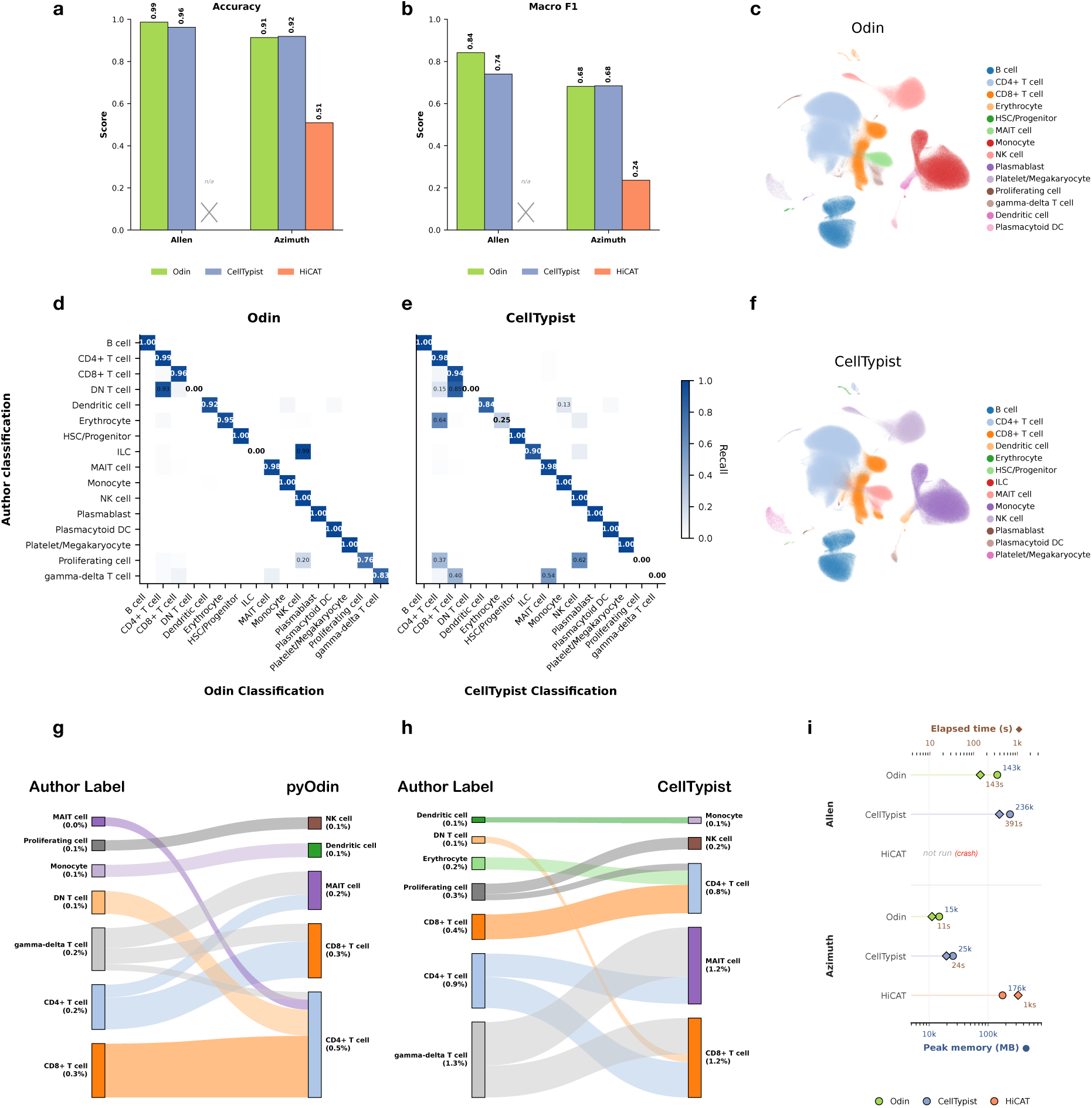
Benchmarking pyODIN against CellTypist and HiCAT for top-level immune cell annotation. **(a)** Overall classification accuracy, defined as the fraction of cells assigned the same label as the author annotation, for pyODIN, CellTypist, and HiCAT across the Allen and Azimuth reference datasets. HiCAT crashed when run on the full Allen dataset due to excessive memory usage (> 512GB) and is therefore only reported for Azimuth. **(b)** Macro-averaged F1-score, calculated as the unweighted mean of the per-class F1-scores and therefore particularly sensitive to performance on rare cell types. **(c)** UMAP of the Allen reference dataset colored by pyODIN top-level cell-type labels. **(d),(e)** Row-normalized confusion matrices for the Allen dataset comparing author annotation (rows) with predicted annotation (columns) for pyODIN **(d)** and CellTypist **(e)**. Diagonal values indicate per-class recall, whereas off-diagonal values indicate the fraction of cells assigned to an alternative label; HiCAT is omitted here as it could not be run on the full Allen dataset. **(f)** UMAP of the Allen reference dataset colored by CellTypist top-level cell-type labels. **(g),(h)** Sankey diagrams summarizing the major misclassification flows (*>* 0.1%) in the Allen dataset for pyODIN **(g)** and CellTypist **(h)**. Ribbons connect author labels (left) to predicted labels (right), with ribbon width proportional to the fraction of alternatively classified cells. **(i)** Peak memory usage (circles, bottom axis) and runtime (diamonds, top axis) for pyODIN, CellTypist, and HiCAT on the Allen and Azimuth datasets, both shown on a log scale.

Because abundant populations can dominate overall accuracy, we next examined the macro-averaged F1-score, which weights each cell type equally and is therefore more sensitive to failures on rare populations. Here, pyODIN again performed strongly, achieving macro-F1 scores of 0.84 and 0.68 on the Allen and Azimuth datasets, respectively, compared with 0.74 and 0.68 for CellTypist and 0.24 for HiCAT on Azimuth (Fig. 2b). These results indicate that pyODIN performs well not only on abundant lineages but also preserves accuracy on rarer, lower-frequency populations, matching or exceeding CellTypist’s performance while substantially outperforming HiCAT.

To understand the structure of these differences, we examined the Allen dataset in more detail. The UMAP of pyODIN-predicted top-level labels showed coherent separation of the major expected immune lineage populations, including B cells, CD4 T cells, CD8 T cells, DCs, monocytes, NK cells, and γδ T cells (Fig. 2c). In contrast, CellTypist failed to recover some populations entirely, most notably γδ T cells (Fig. 2f). Consistent with this, the Allen confusion matrix for pyODIN showed strong concordance with author annotations across most major populations, with only minor confusion between closely related subsets such as ILCs and NK cells (Fig. 2d). CellTypist, by comparison, performed well for abundant populations but showed substantially broader confusion across several rare or difficult-to-distinguish classes (Fig. 2e). Sankey plots of the dominant misclassification flows showed that for pyODIN the most frequent alternative assignments were largely confined to biologically related immune populations (Fig. 2g). CellTypist showed a generally similar overall pattern, although some notable misclassifications were also present, including erythrocyte cells assigned to the CD4 T-cell compartment (Fig. 2h). Together, these results indicate that pyODIN preserves top-level immune structure with high accuracy, exhibiting only limited misassignment between closely related lineages, whereas CellTypist shows broader and more frequent cross-lineage confusion.

The same overall pattern was observed in the Azimuth dataset (Fig. S1). Confusion matrices again showed strong preservation of major populations by pyODIN, whereas CellTypist and HiCAT showed comparatively more misassignment (Fig. S1a-c). Looking at the UMAPs, both pyODIN and CellTypist resolved dissociated clusters of CD8+ cells, whereas HiCAT collapsed them into CD4+ cells (Fig. S1d-f). Furthermore, HiCAT assigned a notable fraction of monocytes to macrophages, a population not typically found in peripheral blood. Sankey plots of the most common alternative labels in Azimuth were similarly more focused for pyODIN, whereas the other methods were more spread out, with HiCAT’s misclassification flows dominated by the monocyte-to-macrophage and CD8+ to CD4+ misclassifications (Fig. S1g-i). These supplementary analyses indicate that the top-level performance patterns observed in Allen generalise across other datasets.

Beyond classification accuracy, computational efficiency is an important practical consideration when annotating large single-cell datasets. We therefore compared peak memory usage and runtime for all three tools across both reference datasets (Fig. 2i). pyODIN was consistently the fastest and most memory-efficient method, annotating the Allen dataset (1,350,748 cells) in 142.7 s using 143 GB of peak memory, compared with 390.9 s and 236 GB for CellTypist. On the smaller Azimuth dataset (161,764 cells), pyODIN required only 11.2 s and 15 GB, again outperforming CellTypist (23.9 s, 25 GB). HiCAT was substantially more resource-intensive, requiring 1,046.8 s and 176 GB of peak memory to annotate the Azimuth dataset alone, consistent with the excessive memory usage that prevented it from being run on the larger Allen dataset. Due to large memory footprint, we thus excluded HiCAT from further analysis and benchmarking. Taken together with the accuracy and macro-F1 results above, these findings indicate that pyODIN combines high classification performance with substantially lower computational overhead relative to existing tools, making it suitable for annotation of large-scale single-cell datasets.

### pyODIN provides finer cell type resolution than CellTypist

We next asked whether pyODIN could better resolve fine-grained cell states than competing annotation tools. To address this, we compared author-curated CD4 T-cell labels with predicted labels from pyODIN and CellTypist. In the Allen dataset, the Sankey plots showed that pyODIN retained substantially greater subtype-level structure than the CellTypist (Fig. 3). pyODIN resolved populations corresponding to naïve, central memory, effector memory, cytotoxic, regulatory, follicular, and helper subsets, including Th1-, Th2-, and Th17-cells (Fig. 3a). By contrast, CellTypist primarily collapsed the CD4 compartment into broader T-cell categories, with more limited subtype specificity and weaker preservation of rare or transitional states (Fig. 3c).

**Figure 3:**
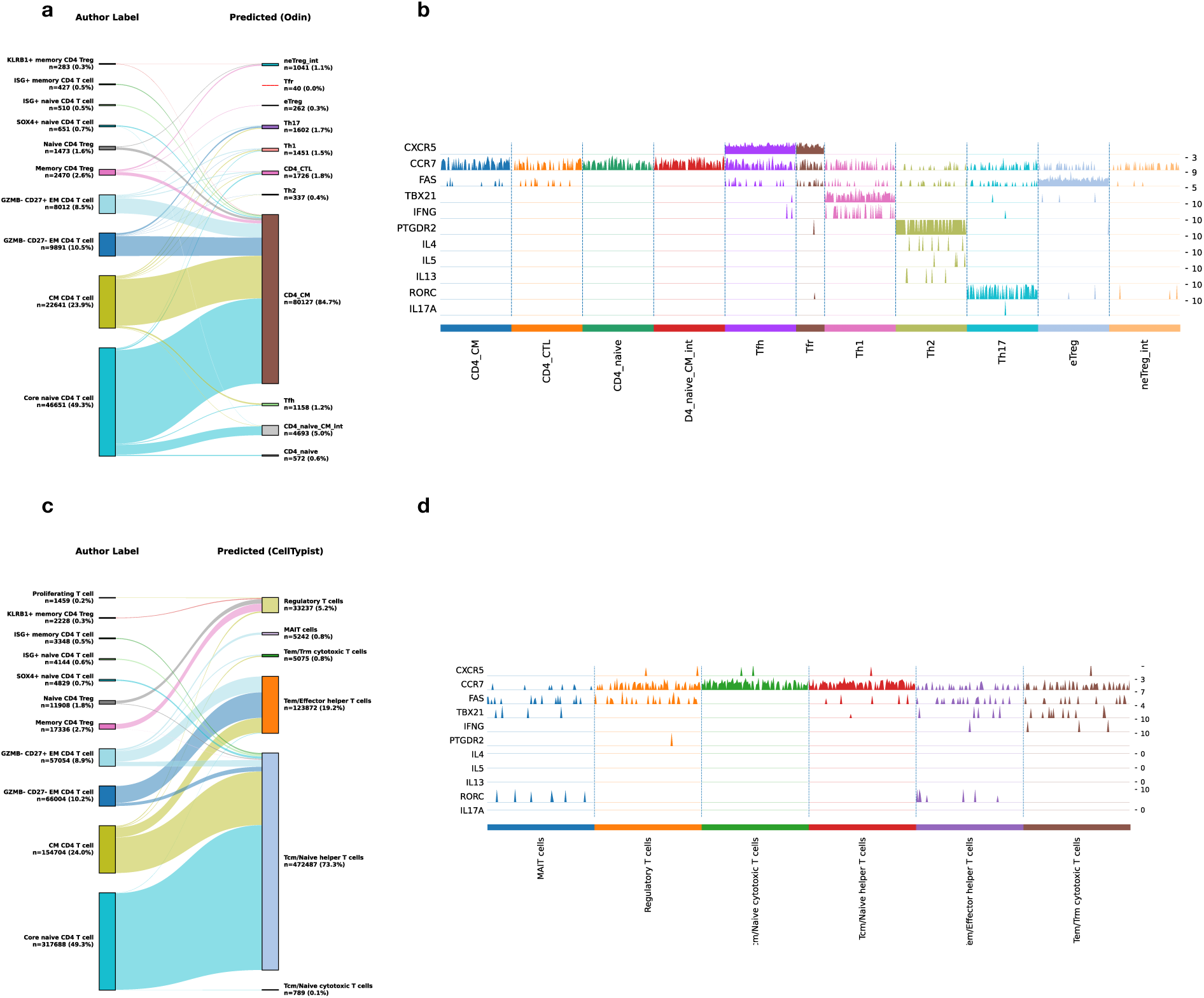
Comparison between pyODIN and CellTypist for fine-grained CD4 T-cell annotation. **(a,c)**, Sankey diagrams showing correspondence between author-curated CD4 T-cell labels (left) and predicted labels (right) for **(a)** pyODIN and **(c)** CellTypist. Ribbon width is proportional to the number of cells assigned to each mapping. **(b,d)**, Marker-gene expression heatmaps across the predicted classes for **(b)** pyODIN and **(d)** CellTypist. Cells are grouped by predicted class, and expression is displayed as scaled values.

The marker-expression heatmaps further supported this difference. In pyODIN, predicted populations were associated with biologically coherent marker patterns, including FOXP3 expression in regulatory subsets such as Treg and Tfr, CXCR5 in follicular populations such as Tfh and Tfr, TBX21 and IFNG in Th1 cells, and CCR7, SELL, CD69, and FAS in populations spanning naïve, central memory, and effector-like phenotypes (Fig. 3b). In contrast, the broader categories returned by CellTypist showed less granular organization of these marker-defined states (Fig. 3d).

A similar pattern was observed in the Azimuth dataset (Fig. S2). pyODIN preserved substantially more subtype-level correspondence to the author-curated CD4 labels than CellTypist, including rare cells such as Tfr, evident in the pyODIN marker-expression heatmap (Fig. S2b). Thus, across both reference atlases, pyODIN retained finer CD4 T-cell structure than CellTypist, which mainly favoured broader T-cell class assignments.

### ADT information rescues lineage assignment when key RNA markers are withheld

We further tested pyODIN against MMoCHi[17] using ADT information from the Azimuth dataset. Like pyODIN, MMoCHi leverages CITE-seq data to assign cell-type annotations. The two tools showed concordant results, though MMoCHi annotations were more granular, since we left its hierarchy unrestricted, whereas pyODIN was limited to top-level annotation (Fig. 4a,b). To directly test the contribution of multimodal information, we ablated the CD8-defining genes CD8A and CD8B from the RNA feature set. Under this condition, pyODIN, relying on RNA-only expression, failed to identify the CD8 T-cell compartment, with cells from that region reassigned to CD4 T cells (Fig. 4c-e). Notably, even

**Figure 4:**
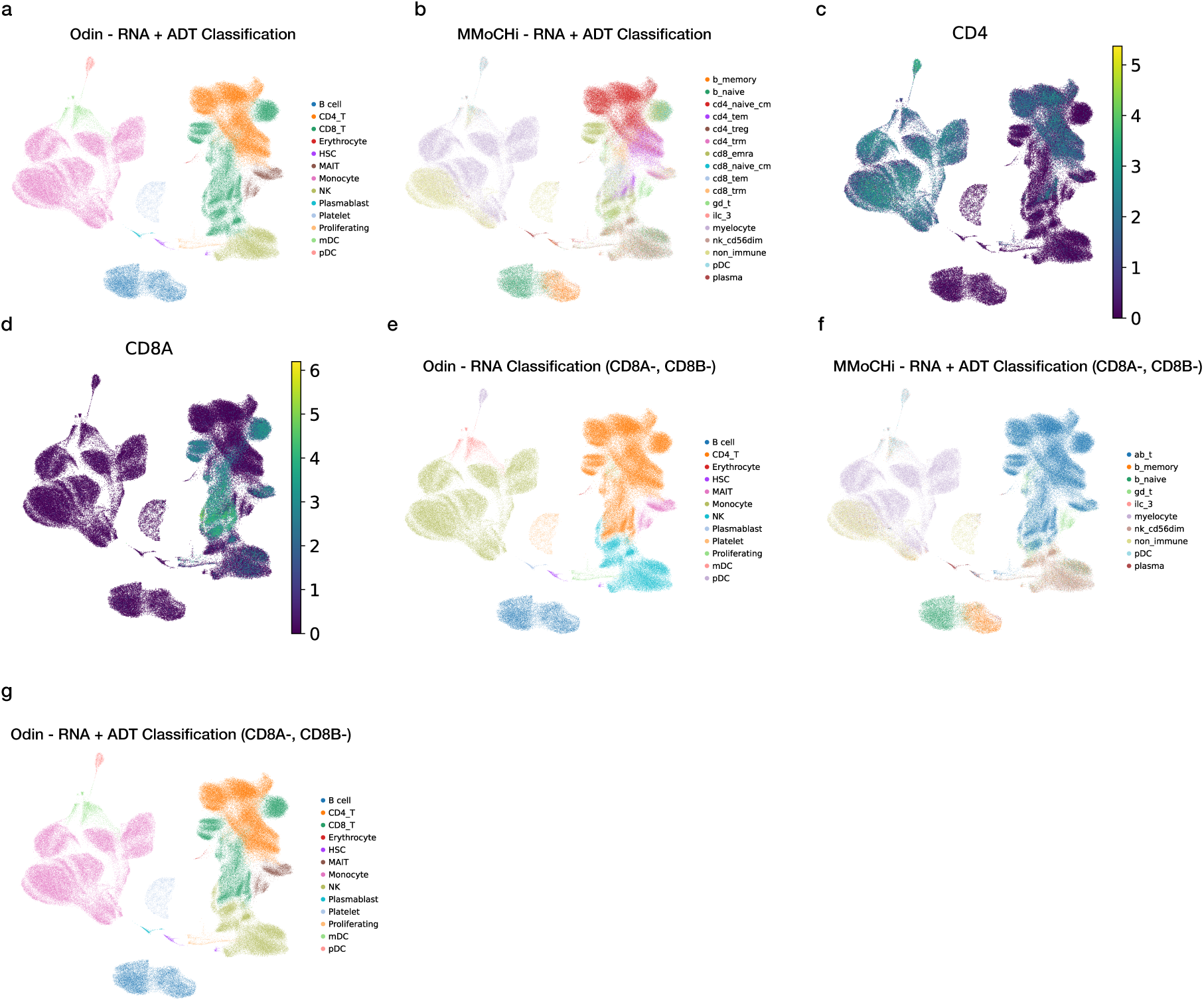
ADT information rescues lineage assignment after ablation of key RNA markers. **(a,b)**,UMAPs showing top-level cell-type annotation by pyODIN **(a)** and MMoCHi **(b)** using combined RNA+ADT input on the Azimuth CITE-seq dataset. Colours indicate predicted cell-type labels; MMoCHi annotations reflect its full unrestricted hierarchy.**c,d**,Feature UMAPs showing expression of CD4 and CD8A. **e-g**, Cell-type annotation after removal of *CD8A* and *CD8B* from the RNA annotation feature set, by pyODIN using RNA-only input **(e)**, MMoCHi using combined RNA+ADT input **(f)**, and pyODIN using combined RNA+ADT input **(g)**. Both pyODIN (RNA-only) and MMoCHi (RNA+ADT) fail to resolve the CD8 T-cell compartment under this ablation, with pyODIN reassigning affected cells to CD4 T cells and MMoCHi collapsing CD4 and CD8 T cells into their shared parent population, αβ T cells. Restoring ADT input to pyODIN **(g)** recovers the CD8 T-cell population despite the same RNA-level ablation.

with RNA and ADT information both present, MMoCHi could not distinguish CD4 and CD8 T cells, collapsing both into αβ T cells - their shared parent population in the MMoCHi hierarchy (Fig. 4f). When ADT information was reintroduced for pyODIN under the same ablation, it recovered the CD8 T-cell population, showing that protein-level evidence can compensate for the loss of corresponding RNA markers (Fig. 4g).

We next tested NK-cell annotation by ablating the NK-associated RNA markers *NCAM1*, *FCGR3A*, and *IL2RB* from the RNA feature set (Fig. S3a-c). Under this condition, MMoCHi collapsed the NK population into the broader nk_ilc supercategory rather than the more precise nk_cd56dim label, even with ADT information present (Fig. S3d). pyODIN, relying on RNA-only expression, failed to identify the NK-cell population, with affected cells reassigned to ILCs (Fig. S3e). When ADT information was reintroduced under the same ablation, pyODIN again recovered the NK-cell population, confirming that protein-level evidence can compensate for the loss of corresponding RNA markers (Fig. S3f).

These two ablation experiments, which independently target CD8 and NK lineage markers, provide a controlled proof of principle for multimodal integration in pyODIN. ADT information acts as a critical fail-safe for lineage assignment when transcriptomic signals alone are insufficient, for example due to RNA dropout events [5]. This kind of refinement is particularly important for distinguishing closely related immune cell subsets, where protein-level markers often provide more robust lineage resolution than gene expression alone.

### Extension of the pyODIN database to support tissue cell types

To support annotation beyond peripheral blood, we extended the pyODIN database to include tissue-associated immune and non-immune cell populations and applied it to liver fine-needle aspirate data[3, 19–22]. At the top level, pyODIN identified broad immune and tissue-associated compartments, including lymphoid populations, monocyte/macrophage compartments, hepatocytes, and liver sinusoidal endothelial cells (LSECs) (Fig. 5a). In subset annotation mode, pyODIN resolved a wider range of immune and tissue-relevant states, including detailed CD4 and CD8 T-cell populations, B-cell subsets, dendritic-cell subsets, macrophage states, and selected non-immune liver cell classes (Fig. 5b). Within the myeloid compartment specifically, pyODIN distinguished fine-grained tissue myeloid subsets, including dendritic-cell and macrophage populations (Fig. 5c), with violin plots of selected marker genes supporting these assignments (Fig. 5d). Expression of TRM-associated markers across CD8 T-cell subsets further supported identification of a CD8 TRM population within the liver dataset (Fig. 5e).

**Figure 5:**
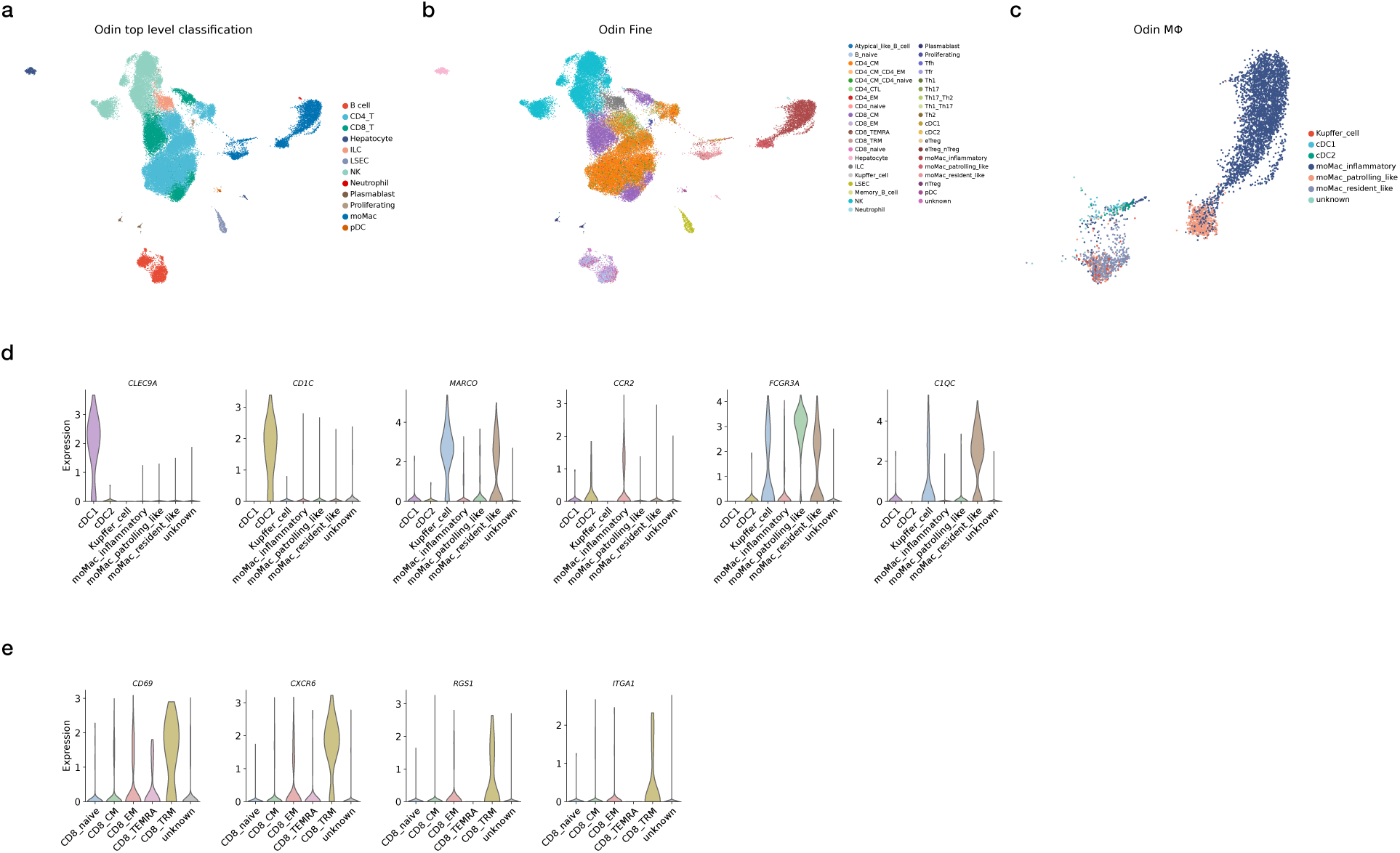
pyODIN enables broad and fine-grained annotation of liver fine-needle aspirate data. **(a)**, Top-level pyODIN annotation of liver fine-needle aspirate single-cell data, showing broad classification of major immune and tissue-associated populations. **(b)**, Fine-grained pyODIN annotation of the same dataset, illustrating higher-resolution assignment of lymphoid, myeloid, macrophage, and tissue-resident populations together with selected non-immune liver cell states. **(c)**, UMAP of the myeloid compartment showing fine-grained pyODIN annotation of tissue subsets. **(d)**, Violin plots showing expression of selected myeloid marker genes across the annotated myeloid populations. **(e)**, Violin plots showing expression of TRM-associated markers across CD8 T-cell subsets.

## Discussion

Accurate immune-cell annotation in single-cell data remains challenging when analyses move beyond broad lineage assignment toward finer immune-state resolution. Closely related immune populations often occupy overlapping transcriptomic space, differ in only a limited number of highly informative markers, and may transition continuously rather than forming sharply separated groups. These challenges are particularly important for rare, transitional, and tissue-adapted states, which are often among the most biologically informative populations but also among the hardest to annotate robustly [1, 4].

pyODIN extends the Odin framework into a broader, more flexible annotation system, combining three features within a single framework: native implementation in Python-centered single-cell workflows, support for multimodal annotation from RNA and ADT data, and expansion of the annotation database from a largely CD4-focused resource into a broader immune and tissue-oriented one. This combination matters because current single-cell studies increasingly span multiple immune lineages, tissue compartments, and data modalities within the same analysis. In addition to these expanded capabilities, pyODIN also runs significantly faster than the original scODIN implementation, an important gain as single-cell datasets continue to grow in scale.

A central contribution of pyODIN is the use of expert-guided immune annotation within a transparent, adaptable framework. In immunology, cell identity is often defined by a small set of lineage-and context-defining markers rather than by global transcriptomic separation alone, particularly for regulatory, helper, follicular, cytotoxic, resident memory, and activated states, where expert interpretation depends on weighting a few highly informative markers. pyODIN makes this logic explicit and easily revisable as subtype definitions evolve [9, 13]. ADT information further improves annotation robustness when key RNA-level lineage markers are weak, missing, or insufficiently discriminative, a scenario likely to be especially useful for closely related lymphoid populations, and in settings where transcript dropout or biological state obscures otherwise canonical markers [6, 7]. The curated marker table may also serve as a practical reference for designing targeted ADT panels for multimodal immune phenotyping when large, high-dimensional panels are unnecessary or impractical.

By expanding the database to include tissue-associated immune and non-immune populations, pyODIN can support annotation beyond PBMC-centered datasets. This is increasingly relevant, as many disease-focused single-cell studies rely on tissue sampling, where immune states must be interpreted alongside parenchymal and stromal compartments. The liver fine-needle aspirate analysis provides an initial demonstration that the framework extends to tissue-relevant annotation settings[9, 12].

This study has certain limitations. Benchmark atlas labels serve as a practical reference standard but do not constitute absolute biological ”ground truth”. Additionally, the multimodal analysis is a targeted proof-of-concept on CD8 T cells and NK cells rather than a comprehensive survey of ADT’s contribution across datasets. Finally, the curated annotation framework demands ongoing expert maintenance, and its performance depends on the quality of the underlying marker definitions. To circumvent this we have implemented a curated gene priority table database with community contributions hosted in the scODIN GitHub repository (https://github.com/jonasns/scodin).

In summary, pyODIN offers a unified annotation framework for single-cell immunology, integrating curated biological knowledge, multimodal evidence, and high-confidence cell classification within a transparent, Python-native workflow. Its principal contribution is enabling immune-cell annotation that jointly addresses broad lineage assignment, fine-grained state resolution, multimodal integration, and cross-tissue extensibility.

## Data availability

The Allen “Sound Life” scRNA-seq data was downloaded from the Allen Institute website (https://apps.allenimmunology.org/aifi/insights/dynamics-imm-health-age/downloads/scrna/, file: SoundLife_OlderAdult_Female_CMVneg.h5ad, 1.35 million cells) and the Azimuth dataset was downloaded using scvi-tools scvi.data.pbmc_seurat_v4_cite_seq() function[23]. The FNA data used in this study are available through dbGaP under accession phs004044.v1.p1 (controlled access).

## Code availability

The pyODIN source code, installation instructions, and documentation are openly available under the MIT License at https://github.com/SondergaardLab/pyODIN. The gene-priority database is provided as supplementary material and will be made publicly available through the repository upon journal acceptance.

## Key Points

• pyODIN is a Python-native, expert-guided framework for cell-type annotation from RNA, antibody-derived tag (ADT), or paired RNA-ADT data, driven entirely by a user-supplied marker database.
• On the Allen and Azimuth immune references, pyODIN matched or exceeded CellTypist and outperformed HiCAT at top level, with fewer cross-lineage errors than either method, and resolved regulatory, follicular helper, memory, and cytotoxic CD4 T-cell states that CellTypist could not resolve.
• On paired RNA-ADT data, pyODIN outperformed MMoCHi. With lineage-defining RNA markers withheld, ADT signal alone recovered CD8 T-cell and NK-cell annotation.
• Extending pyODIN to liver populations required only an expanded marker database and no retraining.
• Vectorised scoring and marker-restricted densification made pyODIN faster and more memory-efficient than the benchmarked alternatives, scaling to more than 1.35 million cells.

## Biographical note

The authors are affiliated with Institut Pasteur de Lille, Université de Lille, and Inserm, and their research spans immunology, metabolic disease, single-cell biology, and computational bioinformatics

## Acknowledgment

This project is funded by the European Union - European Regional Development Fund (FEDER 25000736/HDF008084), Métropole Européenne de Lille (MEL), and Institut Pasteur de Lille to JNS. JTH is supported by an ERC Starting Grant (Metabo3DC, contract number 101042759). DD is supported by grants from the Agence Nationale de la Recherche (ANR) and the European Union: EGID ANR-10-LABX-46 and ANR-18-CE15-0024, ANR-20-CE15-0026, ANR-21-CE15-0020, ANR-23-CE15-0032-01), Fondation pour la Recherche Médicale (FRM).

This work used data from the liver fine-needle aspirate single-cell RNA-seq component of the “Single-cell atlas of human liver and blood immune cells across fatty liver disease stages” study (dbGaP accession phs004044.v1.p1). The study was conducted by the original investigators and supported by the National Institute of Diabetes and Digestive and Kidney Diseases (NIDDK). The data were supplied by Dr. Nadia Alatrakchi and Massachusetts General Hospital. This manuscript was not prepared in collaboration with the investigators of that study and does not necessarily reflect their views or those of NIDDK.

## Author contribution

Conceptualization: JNS. Methodology: SST and JNS. Software: SST. Data curation: JTH, BS, DD, and JNS. Formal analysis: SST and JNS. Visualization: SST and JNS. Supervision: JNS. Project administration: JNS. Funding acquisition: JNS. Writing - original draft: SST and JNS. Writing - review and editing: all authors.

## Supplementary Material

### Supplementary Figures

**Figure S1:**
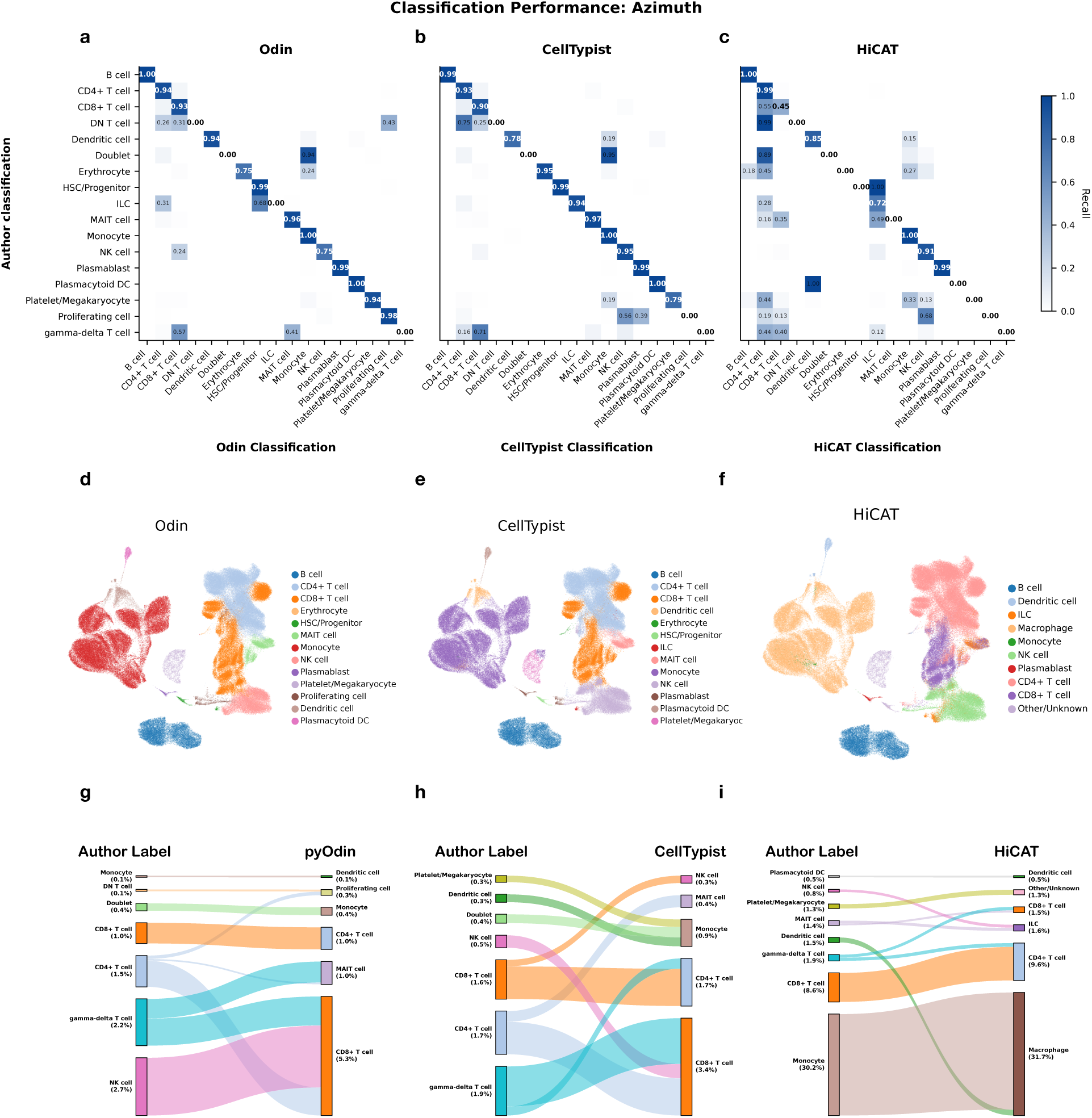
Classification performance on the Azimuth dataset. (a)-(c) Row-normalized confusion matrices for the Azimuth dataset comparing author annotation (rows) with predicted annotation (columns) for pyODIN **(a)**, CellTypist **(b)**, and HiCAT **(c)**. Diagonal values indicate per-class recall, whereas off-diagonal values indicate the fraction of cells assigned to an alternative label. **(d)-(f)** UMAP of the Azimuth reference dataset colored by top-level cell-type labels predicted by pyODIN **(d)**, CellTypist **(e)**, and HiCAT **(f)**. **(g)-(i)** Sankey diagrams summarizing the major misclassification flows in the Azimuth dataset for pyODIN **(g)**, CellTypist **(h)**, and HiCAT **(i)**. Ribbons connect author labels (left) to predicted labels (right), with ribbon width proportional to the fraction of alternatively classified cells.

**Figure S2:**
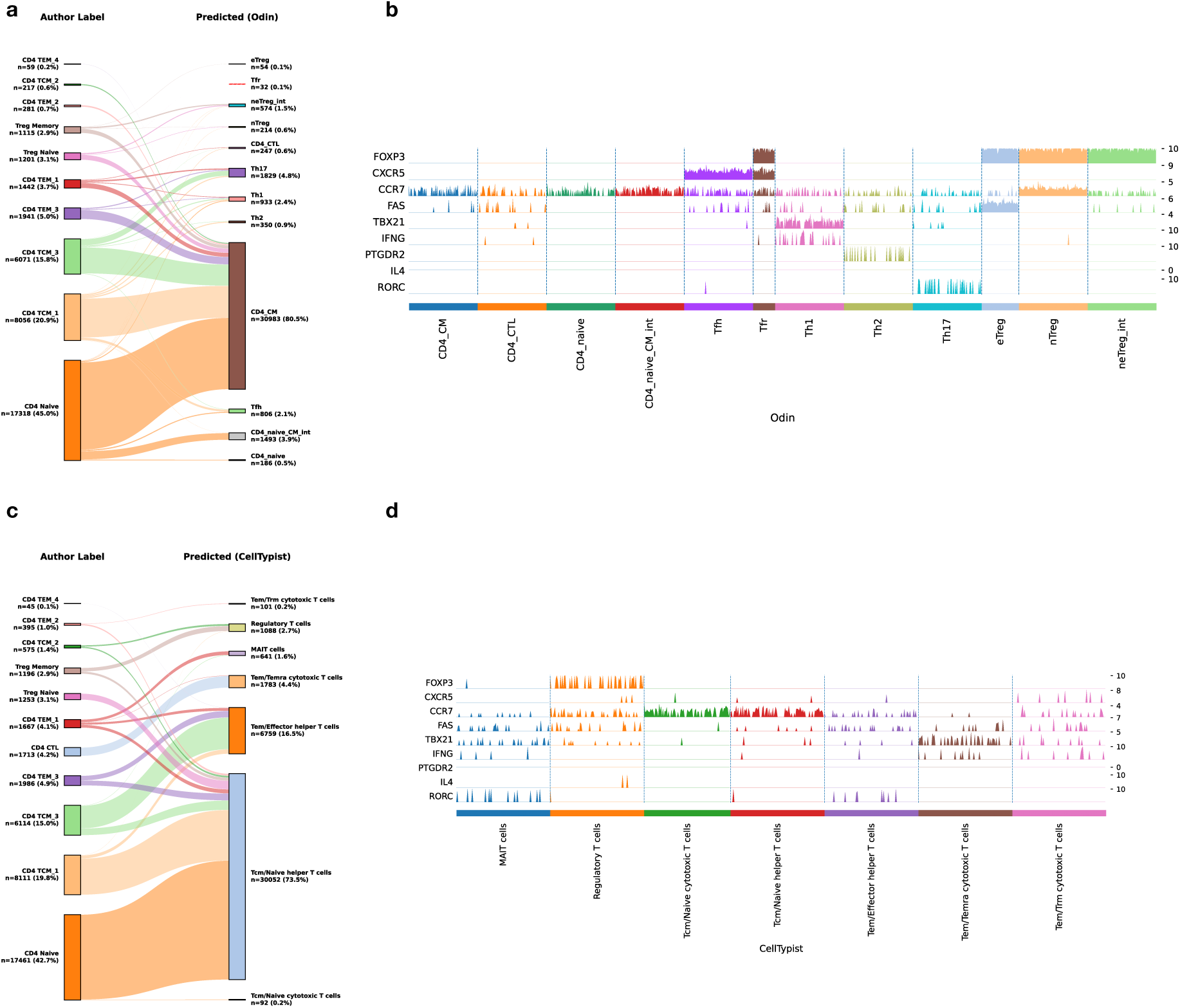
Fine-grained CD4 T-cell annotation in the Azimuth reference dataset. **(a,c)**, Sankey diagrams showing correspondence between author-curated CD4 T-cell labels (left) and predicted labels (right) for **(a)** pyODIN and **(c)** CellTypist in the Azimuth dataset. Ribbon width is proportional to the number of cells assigned to each mapping. **(b,d)**, Marker-gene expression heatmaps across the predicted classes for **(b)** pyODIN and **(d)** CellTypist. Cells are grouped by predicted class, and expression is displayed as scaled values.

**Figure S3:**
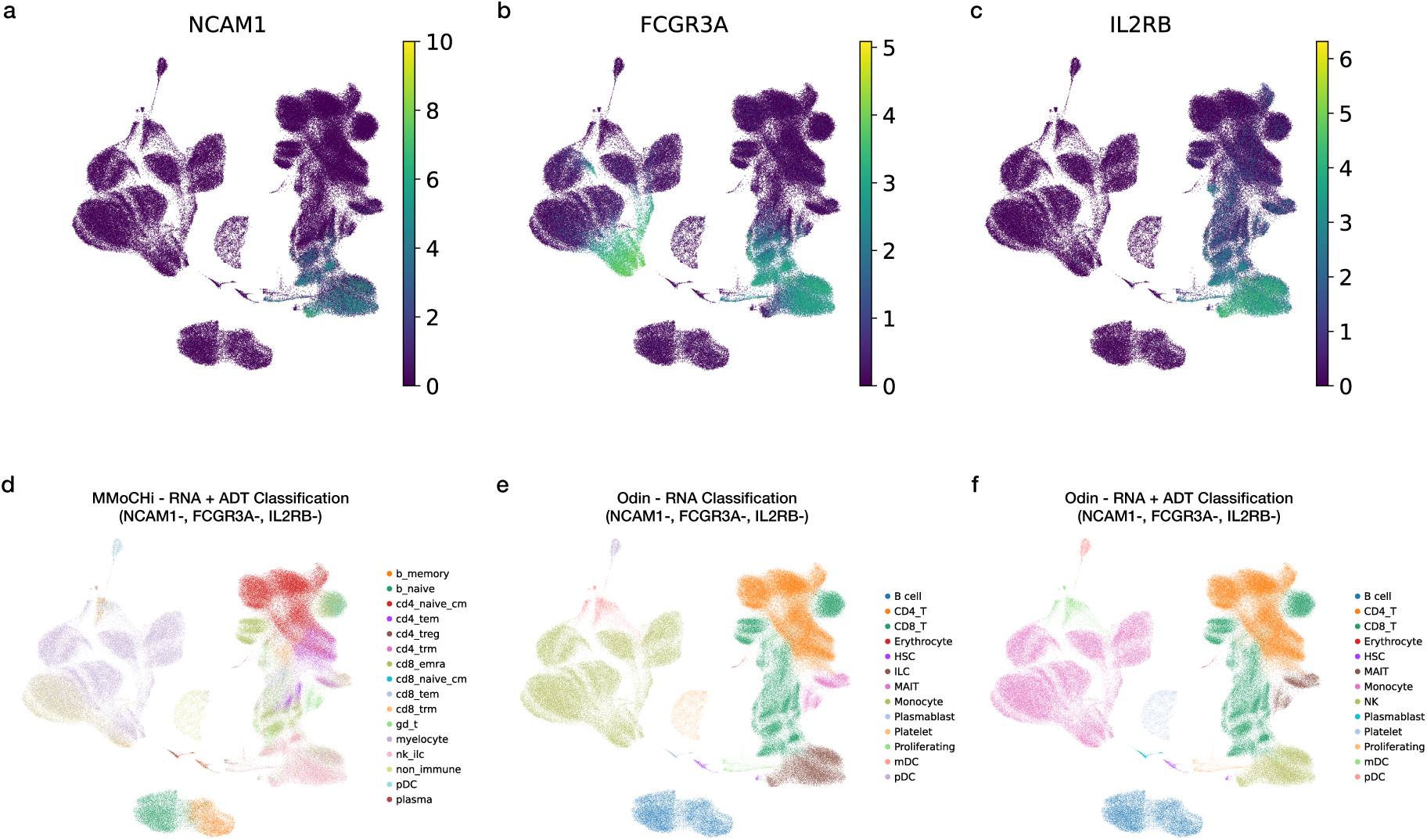
ADT information rescues NK cell identification. **(a-c)**, Feature UMAPs showing expression of NK lineage markers *NCAM1*, *FCGR3A*, and *IL2RB*. **(d-f)**, Cell-type annotation after removal of *NCAM1*, *FCGR3A*, and *IL2RB* from the RNA annotation feature set, by MMoCHi using combined RNA+ADT input **(d)**, pyODIN using RNA-only input **(e)**, and pyODIN using combined RNA+ADT input **(f)**. MMoCHi collapses the NK population into the broader nk_ilc supercategory rather than the more precise nk_cd56dim label, despite ADT information being present **(d)**. pyODIN using RNA-only input fails to identify the NK-cell population, with affected cells reassigned to ILCs **(e)**. Restoring ADT input to pyODIN **(f)** recovers the NK-cell population despite the same RNA-level ablation.

### Supplementary Tables

**Table S1:**
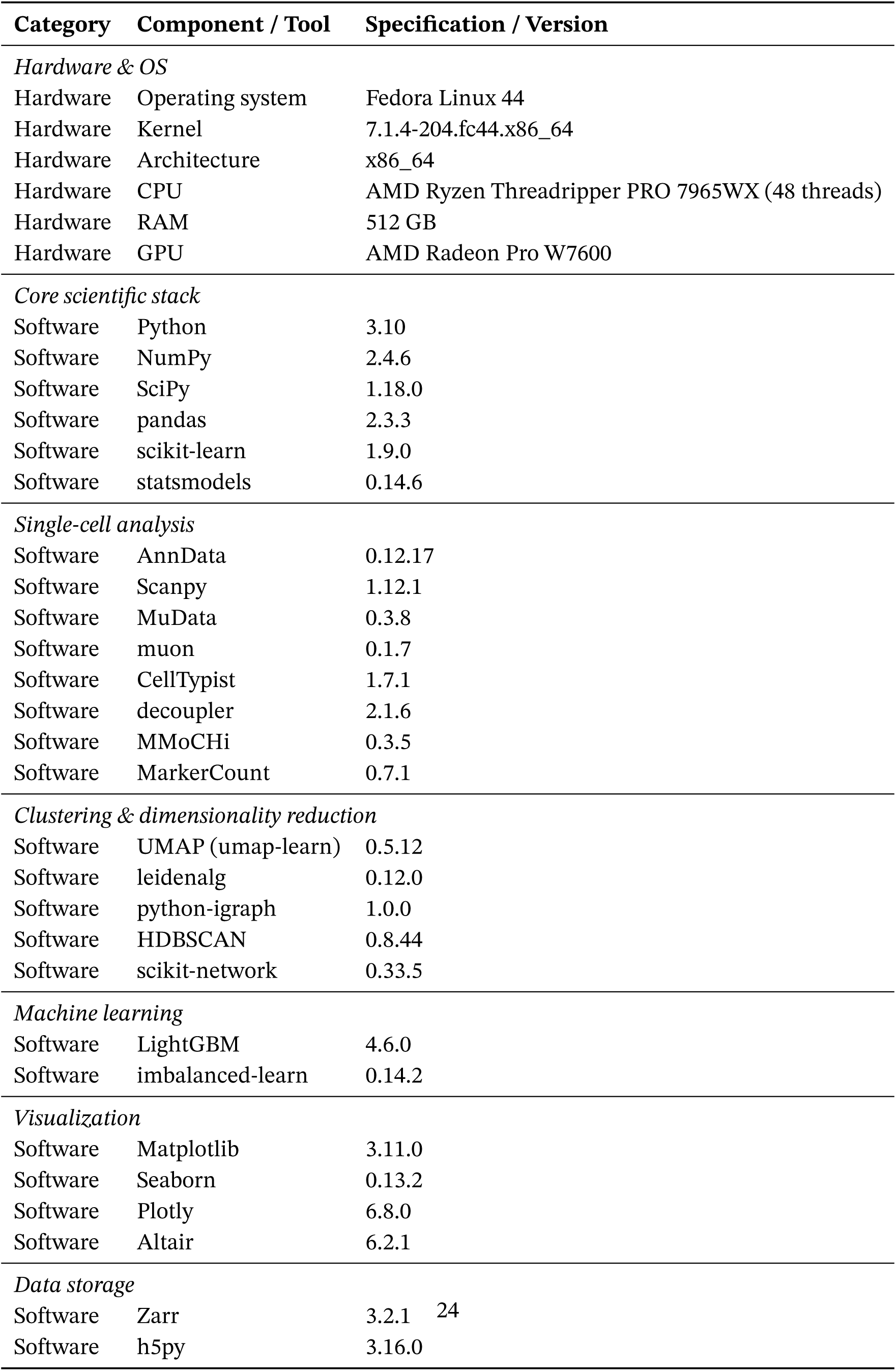
Software and hardware specifications used in this manuscript.

**Table S2:** Cell label harmonization table.

| Harmonized Category | Author | CellTypist | HiCAT | Odin |
| --- | --- | --- | --- | --- |
| B cell | B intermediate, B memory, B naive, Effector B cell, Memory B cell, Naive B cell, Transitional B cell | Age-associated B cells, B cells, Memory B cells, Naive B cells, Proliferative germinal center B cells, Transitional B cells | B cell | B cell |
| CD4+ T cell | CD4 CTL, CD4 Naive, CD4 TCM, CD4 TEM, Memory CD4 T cell, Naive CD4 T cell, Treg | Follicular helper T cells, Memory CD4+ cytotoxic T cells, Regulatory T cells, T(agonist), Tcm/Naive helper T cells, Tem/Effector helper T cells, Treg(diff), Type 1 helper T cells, Type 17 helper T cells | T cell CD4+ | CD4_T |
| CD8+ T cell | CD8 Naive, CD8 TCM, CD8 TEM, CD8aa, Memory CD8 T cell, Naive CD8 T cell | CD8a/a, CD8a/b(entry), Tcm/Naive cytotoxic T cells, Tem/Temra cytotoxic T cells, Tem/Trm cytotoxic T cells, Trm cytotoxic T cells | T cell CD8+ | CD8_T |
| DN T cell | DN T cell, dnT | Double-negative thymocytes | - | - |
| Dendritic cell | ASDC, cDC1, cDC2 | DC precursor, DC1, DC2, DC3 | Dendritic cell | mDC |
| Doublet | Doublet | - | - | - |
| Erythrocyte | Eryth, Erythrocyte | Early erythroid, Late erythroid, Mid erythroid | - | Erythrocyte |
| HSC/Progenitor | HSPC, Progenitor cell | ELP, ETP, Early lymphoid/T lymphoid, HSC/MPP, Megakaryocyte-erythroid-mast cell progenitor, Myelocytes | - | HSC |
| ILC | ILC | ILC3 | ILC | ILC |
| MAIT cell | MAIT | MAIT cells | - | MAIT |
| Macrophage | - | Kupffer cells, Macrophages | Macrophage | - |
| Mast cell | - | Mast cells | - | - |
| Monocyte | CD14 Mono, CD14 monocyte, CD16 Mono, CD16 monocyte, Intermediate monocyte | Classical monocytes, Monomac, Monocyte precursor, Monocytes, Non-classical monocytes | Monocyte | Monocyte |
| NK cell | CD56bright NK cell, CD56dim NK cell, NK, NK_CD56bright | CD16+ NK cells, CD16- NK cells, NK cells | NK cell | NK |
| NKT cell | - | NKT cells | - | - |
| Other/Unknown | - | Double-positive thymocytes | unassigned | - |
| Plasmablast | Plasma cell, Plasmablast | Plasma cells, Plasmablasts | Plasma cell | Plasmablast |
| Plasmacytoid DC | pDC | pDC | - | pDC |
| Platelet/Megakaryocyte | Platelet | Megakaryocyte precursor, Megakaryocytes/platelets | - | Platelet |
| Proliferating cell | CD4 Proliferating, CD8 Proliferating, NK Proliferating, Proliferating NK cell, Proliferating T cell | Cycling T cells | - | Proliferating |
| gamma-delta T cell | gdT | CRTAM+ gamma-delta T cells | - | gdT |

## Notes

### Competing Interest Statement

The authors have declared no competing interest.

### Summary of Updates

The manuscript has been substantially revised and expanded. Benchmarking now includes a larger part of the Allen immune dataset (1.35 million cells). We also added comparisons with MMoCHi, together with controlled marker ablation experiments. Runtime and peak memory usage were systematically compared across pyODIN, CellTypist, and HiCAT. The methods section has been expanded to describe the vectorised scoring framework, hierarchical annotation logic, cluster consensus procedure, LightGBM based classification of unresolved cells, and memory efficient implementation. Figures, supplementary analyses, the abstract, title, discussion, documentation, and code availability information have been updated accordingly.

